# Automated counting of *Drosophila* imaginal disc cell nuclei

**DOI:** 10.1101/2023.05.26.542420

**Authors:** Pablo Sanchez Bosch, Jeffrey D. Axelrod

## Abstract

Automated image quantification workflows have dramatically improved over the past decade, enriching image analysis and enhancing the ability to achieve statistical power. These analyses have proved especially useful for studies in organisms such as *Drosophila melanogaster*, where it is relatively simple to obtain high sample numbers for downstream analyses. However, the developing wing, an intensively utilized structure in developmental biology, has eluded efficient cell counting workflows due to its highly dense cellular population. Here, we present efficient automated cell counting workflows capable of quantifying cells in the developing wing. Our workflows can count the total number of cells or count cells in clones labeled with a fluorescent nuclear marker in imaginal discs. Moreover, by training a machine-learning algorithm we have developed a workflow capable of segmenting and counting twin-spot labeled nuclei, a challenging problem requiring distinguishing heterozygous and homozygous cells in a background of regionally varying intensity. Our workflows could potentially be applied to any tissue with high cellular density, as they are structure-agnostic, and only require a nuclear label to segment and count cells.

## Introduction

*Drosophila* imaginal discs have been an important and productive model for studying cellular and molecular processes for several decades. Of the multiple fly imaginal discs, the most widely utilized is the wing disc. The wing disc starts as a primordium of about 50 cells at the conclusion of embryonic development. By the end of the larval development, it is populated by about 30.000 to 40.000 cells (Garcia-Bellido and Merriam, 1971; Martín *et al*., 2009). This massive expansion in cell number takes place during only about 96 hours. Therefore, the wing disc provides an excellent *in vivo* model to study processes that regulate cell growth and survival, and the molecular mechanisms that protect multicellular organisms from deleterious and proto-oncogenic mutations that modify cell fitness (Held Jr, 2005; Baena-Lopez *et al*., 2012). Using mosaic analysis (Xu and Rubin, 1993; Perrimon, 1998; Lai and Lee, 2006), investigators can assess the effect of any gene on cell proliferation, death, and overall fitness just by comparing the relative growth of a mosaic clone to the surrounding wildtype (WT) environment over the course of disc development (Perrimon, 1998; Rogulja and Irvine, 2005; Martín *et al*., 2009; Amoyel and Bach, 2014). These characteristics of imaginal discs have enabled many important contributions to our understanding of the mechanisms regulating cell proliferation, cell death, tumorigenesis, and cell competition (Brumby and Richardson, 2003; de la Cova *et al*., 2004; Johnston, 2014; Ferreira and Milán, 2015).

Although the wing disc provides an excellent system to study these processes, the nature of disc structure substantially limits the accuracy of typically applied methods to quantify disc or clone growth in terms of cell numbers. Cell counting techniques usually rely on nuclear or membrane staining, and despite being a simple columnar epithelium, the densely populated disc proper and the folds that form during growth pose a problem for image analysis workflows that would seek to directly count cells. For this reason, researchers have typically quantified cell proliferation, death, and cell competition by either measuring the area covered by the cells of the analyzed clones as a surrogate for cell number, or by manual counts. Area measurements are based on the assumption that all cells occupy a similar surface area, i.e. two clones of similar cell count will cover the same surface area of the imaginal disc. While permitting a rapid manual analysis on a large number of clones, it produces substantial errors for several scenarios in which the similar surface area assumption is wrong. Clones of similar cell number cover very different surface areas depending on the region of the disc due to the variation in cell compaction and due to the folds that naturally form during disc maturation (Day and Lawrence, 2000) and Fig 1B). Furthermore, clones carrying mutations that alter structural characteristics of cells can disproportionately modify cell number and the occupied surface area; in these situations, it would be advantageous to quantify these effects independently. Therefore, directly counting the number of cells in a clone and the total cell count in the imaginal disc would provide a more accurate assessment of tissue growth dynamics compared to typically used methods. The large numbers of cells involved in these experiments makes manual counting a laborious and time-consuming task, limiting the number of biological replicates that can be included in statistical analyses. Given recent advances in image processing and quantification, we explored the possibility of automating the counting of labeled nuclei in wing imaginal discs and in clones within discs. Here, we provide effective protocols based on readily available open-source image analysis software and plugins, that enable fast and accurate quantification of cell counts in the wing imaginal disc. Moreover, our protocol can be easily adapted to other discs and similar epithelial structures where dense and variable cell packing and large numbers would challenge other methods.

**Figure 1.**
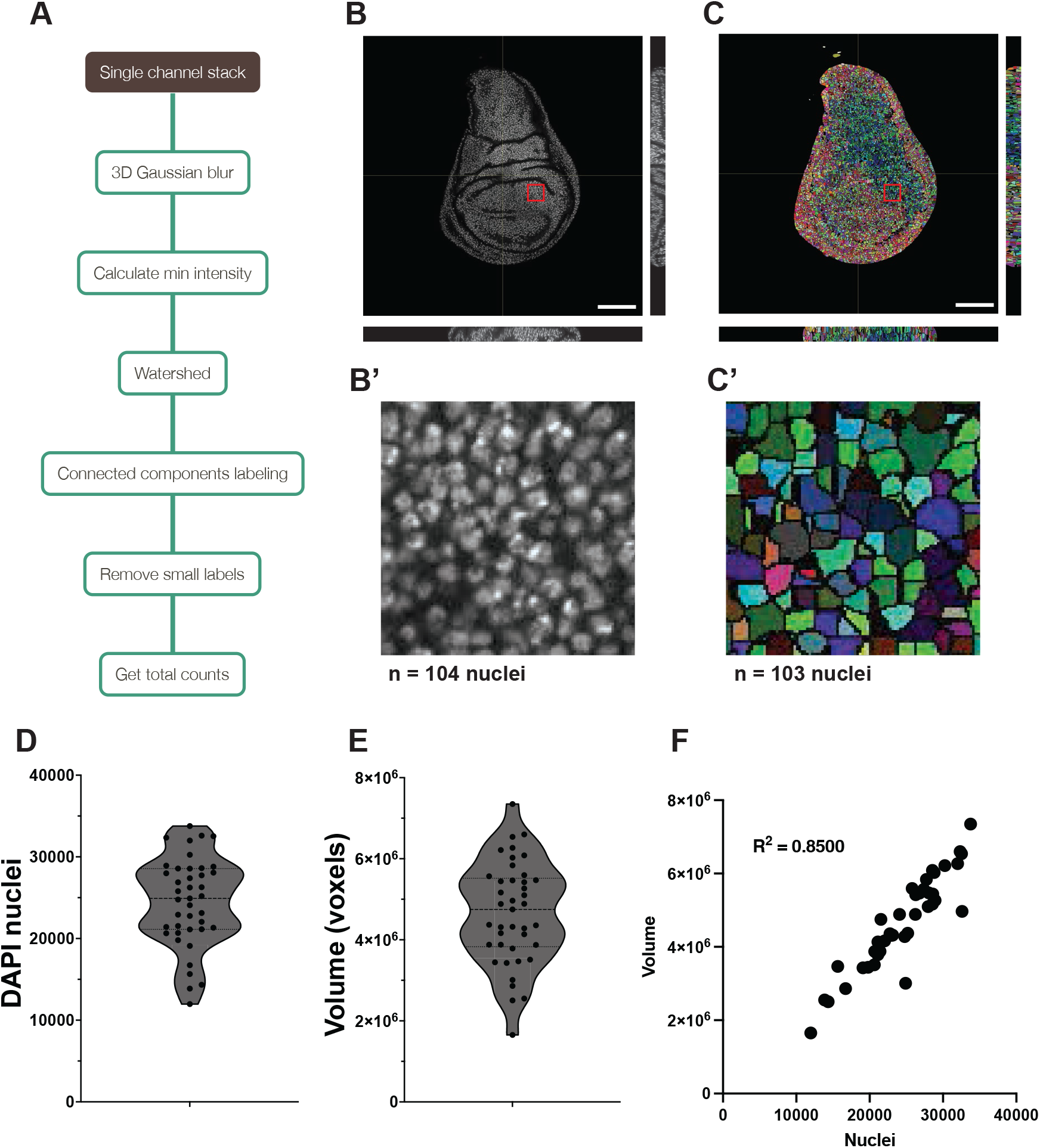
Automated counting of DAPI nuclei. A) Schematic of the counting workflow. B) Overview of DAPI-stained nuclei with orthogonal views to show the overall nuclear density. C) Watershed of the wing disc shown in B, with each nucleus in a different color. B’ and C’) show a close-up (indicated by a square in B and C) of the wing pouch. Manual counting of the close-up window is shown below the images. D) Violin plot showing the individual nuclear counts, mean, and standard deviation of 40 wing discs. E) Violin plot showing the individual disc volume, median and standard deviation of the same 40 wing discs. F) Correlation plot of nuclear counts vs. volume of the 40 discs, showing an R^2^ of 0.8112. Scale bar: 100 μm

## Results and Discussion

Imaginal disc nuclei are densely packed and commonly overlap when viewed in X-Y plane images of the pseudostratified epithelium. To accurately separate and count nuclei in the wing imaginal disc we have designed a workflow that performs two main steps:

1. Segmentation of nuclei in 3D via watershedding
2. Object classification and counting

We first describe a method using these two simple steps to obtain the cell counts of both DAPI-stained nuclei and clones marked with a nuclear fluorescent tag.

We next describe an expansion of our protocol to enable quantification of cells from homozygous clones in twin-spot analysis. Twin-spots are produced by the mitotic recombination of a chromosomal arm. In the simplest form of twin-spot analysis, one of the two chromosome arms carries a fluorescent marker (usually eGFP or RFP). After mitotic recombination, the clones generated will carry either two copies of the fluorescent marker, or no marker at all. Thus, the resulting tissue will have three distinct populations:

- Heterozygous, non-recombinant cells
- Homozygous, fluorescent recombinant cells
- Homozygous non-marked recombinant cells

The two homozygous populations are generated from a single cell and under wildtype conditions will grow into twin clones containing roughly the same number of cells. The existence of a background of heterozygous cells makes it impossible to apply the same workflow used for marked clones in an unmarked background, as it will also count the heterozygous cells. Recent advances in image analysis and trainable ML algorithms allowed us to create a workflow that separates homozygous from heterozygous cells for automated twin-spot cell counts. By adding an ML-trainable segmentation to our cell count analysis, we achieved automatic classification of homozygous twin clones, distinguishing them from heterozygous non-recombinant cells. After the segmentation of twin-spots, we can count the nuclei inside the clone and its twin-spot to obtain a reliable readout of cellular processes affecting cell division and survival (An overview of all three workflows is outlined in Suppl. Fig. 1).

### Segmenting and counting DAPI-stained nuclei

Accurate counting of cell numbers is essential for quantitative representation of tissue growth. Counting the total number of cells in *Drosophila* wing imaginal discs has been extremely challenging, as the pseudostratified epithelium is densely packed with narrow cells and nuclei are very close to each other and overlap along the Z axis (apical-basal orientation) [REF]. The most thorough quantification of cells in the wing disc as of the publication of this work estimates that at the end of larval development the disc contains around 31.000 cells(Martín *et al*., 2009), although other publications set this number around 35.000 and up to 50.000 cells. Wing disc development can vary between individuals and two larvae may have very disparate cell numbers in their wing discs. This prompted us to find a method to score and count all cells in the wing imaginal disc quickly and accurately.

To count wing disc nuclei, we follow a sequence of steps (Fig. 1A) beginning with a simple DAPI staining to acquire a high-quality Z stack of the wing imaginal disc as detailed in our methods (Fig. 1B). Starting from a high-quality image is crucial for proper counting. We loaded the images into Fiji and, if present, we cropped out the haltere and leg discs. Last, we ran the nuclear count workflow macro (“DAPI_counts_with_MorphoLibJ.ijm”; Fig. 1A and described in the methods) to count DAPI-stained nuclei in Fiji via Watershed and Connected Components Labeling (Fig. 1C). We performed the analysis on 40 discs.

The step in this protocol with the largest influence on the final counts is the noise removal and smoothing performed via a 3D Gaussian blur. For our images, we found that a 2 px 3D Gaussian blur rendered the best results for the downstream analysis (Suppl. Fig 2). For quality check, we manually counted smaller regions of the wing disc to ensure the watershed segmentation worked as intended (Fig. 1B’ and C’). Depending on the image quality, this parameter might need adjustment, although all our tests suggested that the majority of images perform best with a 2 px Gaussian Blur.

Our analysis shows a high variability of cell counts in third instar larval discs, ranging from 20.000 to 35.000 cells (Fig. 1D). While it may be that this range reflects significant differences in the cell numbers of mature wing discs between individuals, some of this range may be accounted for by the possibility that discs are harvested before the last cell divisions occur. We further analyzed the correlation of total nuclei with disc volume and area. Disc volume was somewhat less variable than cell counts, with fewer outlier discs and most samples falling closer to the average volume (Fig. 1E). Although the number of nuclei correlates with disc volume (Fig. 1F), disc volume is not a robust surrogate for cell counts, having a correlation coefficient of R^2^ = 0.85. Furthermore, disc areas measured from the max projection of the disc stack are even worse proxies of cell count. Areas poorly correlate with both total volume occupied (Suppl. Fig. 3A, R^2^ = 0.5851) and nuclear counts (Suppl. Fig. 3B, R^2^ = 0.59).

Several attempts have been made over the decades to estimate the number of cells in the wing disc at the end of larval development, showing very disparate final counts ranging between 30.000 and 50.000 cells (Milán *et al*., 1996; McClure and Schubiger, 2005; Martín *et al*., 2009). The most recent estimations, based on manual counting, set the number between 30.000 and 40.000 cells (McClure and Schubiger, 2005; Martín *et al*., 2009). Our results agree with these more recent estimates (McClure and Schubiger, 2005) and manual nuclear counts (Martín *et al*., 2009). Our analysis is to date the only fast and accurate measure of wing disc cell number and volume.

### Segmenting and counting fluorescently tagged clones in imaginal discs

Total cell counts can be applicable to many studies, but the most common scenario in which investigators wish to quantify cell number is in the assessment of clone sizes. Generating fluorescently tagged clones that harbor a mutation in the wing disc is a widely used method to study tissue growth, programmed cell death, cell competition, or tumorigenesis. In the cases in which mutations have the most dramatic effects, simple observation of clone sizes compared to twin-spots or control clones may be sufficient to qualitatively determine the effect of the mutation. However, mutations inducing weaker though potentially important effects, or those that may separately modify cellular structure and cell number (e.g. cadherins, components of the apical junctions, etc (Lecuit *et al*., 2011; Borghi *et al*., 2012; Fulford and McNeill, 2020)) will require quantitative measurement of cell number and volume of clones to appropriately describe their effects.

Here we show that the workflow used to count DAPI-stained nuclei in the whole disc is similarly effective in counting cell numbers in wing disc clones expressing a fluorescent protein fused to the Nuclear Localization Sequence (NLS). A simple expansion of the workflow enables it to also register the total number of clones and the average cell count per clone, which can provide additional information in processes such as the elimination of cells and clones by cell competition.

We created clones in the wing disc tagged with either NLS::GFP or NLS::RFP in a WT background to test our workflow. Fig. 2A shows one of the analyzed discs with NLS::GFP cells, with the corresponding segmentation shown in Fig. 2B. For quality control we manually counted cells in five clones and validated nearly perfect counting by our workflow (Fig. 2C and C’ and Fig 2D and D’).

**Figure 2.**
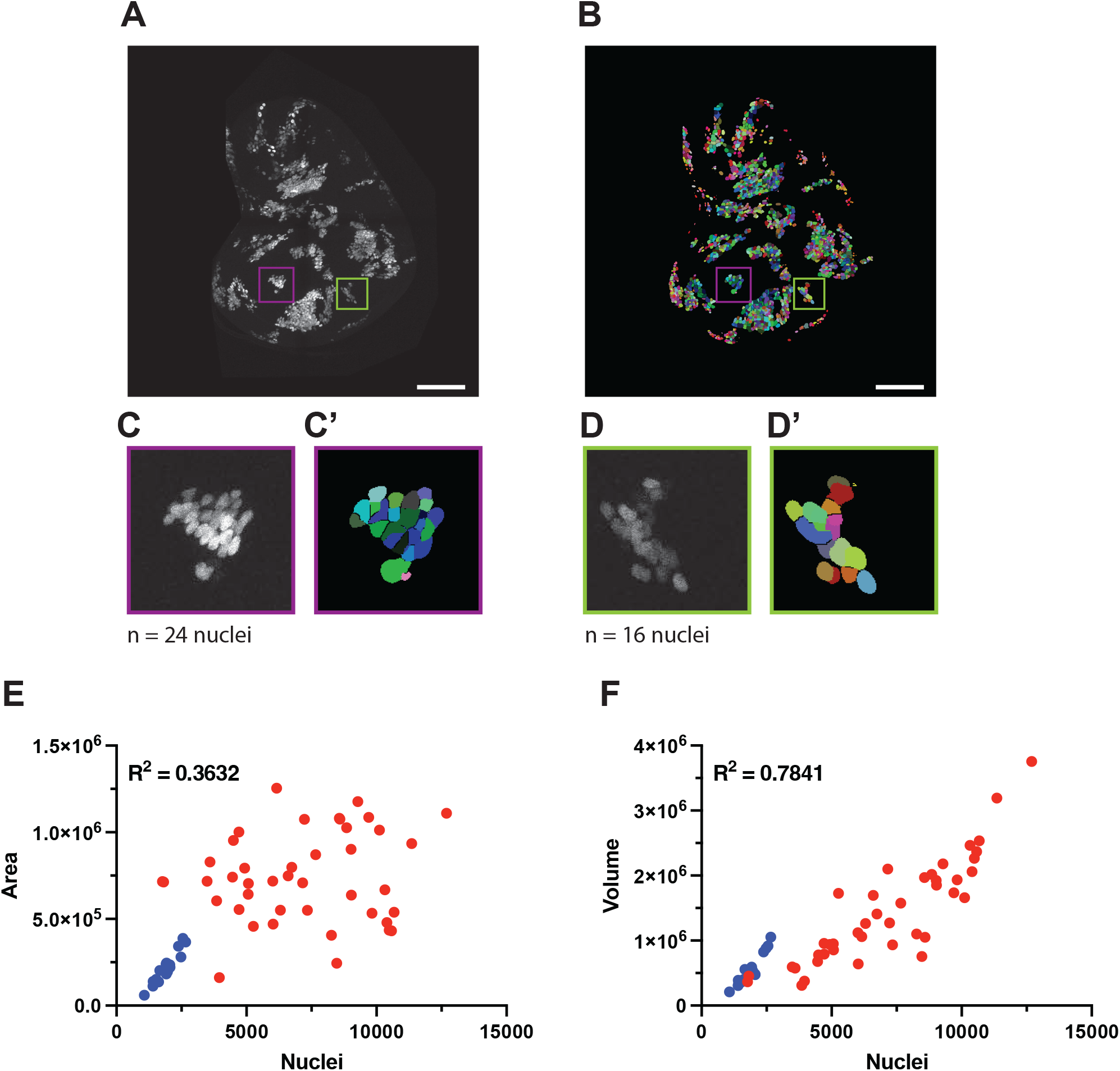
Automated counting of nGFP-marked clones. A) Max projection of the Z-stack of a representative wing disc with nGFP clones. B) Max projection of the Watershed of the disc shown in A. C and C’) Closeups of the clone highlighted in purple in A and B. Manual count of nuclei, shown below the image, returns the same number for both the original image and the watershed. D And D’) Closeups of the clone highlighted in green in A and B. Manual count of nuclei, shown below the image, also returns the same number for both the original image and the watershed. A total of 5 clones were manually counted. The pipeline miscounted 1 or 2 cells in clones of 100 or more cells, and made no errors for smaller clones. E) Correlation between nuclear counts and area in the whole dataset, comprising 17 discs tagged with NLS::GFP (blue) and 42 discs marked with NLS::RFP (red). F) Correlation between nuclear counts and volume in the same dataset of 17 NLS::GFP and 42 NLS::RFP. Scale bar: 100 μm

Typically, wing disc clones have been quantified by measuring the area the clones occupy in the disc. However, we have already shown that areas, unlike volumes, are a poor proxy for cell size for whole discs (Fig. 1F and Suppl. Fig. 3). We decided to evaluate the correlation between cell counts, area, and volume for wing disc clones. To do so, we counted two datasets, comprised of 17 discs tagged with NLS::GFP with lower density of clones, and 42 discs tagged with NLS::RFP with higher clonal density. Similar to our observations with DAPI nuclei, the area covered by the clone is a poor replacement for total counts (R^2^ = 0.3632; Fig. 2E). The total volume of clonal cells shows a somewhat better correlation with total counts (R^2^ = 0.7841; Fig. 2F). If we take the lower clonal density NLS::GFP dataset alone, both area and volume provide a better readout of the clone size (Suppl. Fig. 4, R^2^ = 0.9107 for area and R^2^ = 0.8922 for volume). As discussed above, the poor correlation between cell number and area may be due to regional variation in cell compaction and the folding of the epithelium that occurs naturally during disc maturation. Importantly, however, the poor performance of area as a proxy for cell number is worse at higher clone density. Therefore, studies that use clone area as a proxy for cell number should be interpreted with caution. On the other hand, the relatively stronger correlation between cell number and clone volume suggests a relatively uniform cell volume across the disc, and provides an opportunity to add an additional dimension to clonal analyses. In wildtype clones, clone volumes correlate to cell numbers, whereas in some mutant clones, a discordance between cell number and clone volume compared to wildtype tissue could reveal that the mutant induces diminished or excess cell size, thus providing a more nuanced readout of the phenotypic effects of a given mutant on the wing disc cells.

In some instances, measuring the number of clones per disc and the average cell number per clone might prove useful. We extended our counting workflow to also quantify these values. To do so, we simply merged neighboring nuclear watersheds using Fiji’s Dilate function and then performed the same segmentation and counting method used for cell counting, which in this case counts clones made of fused neighbors. Finally, we isolated each clone and counted the nuclei within its boundaries to obtain the average clone size (Fig. 3A-D, full description in the methods section). To avoid counting as separate clones isolated single marked cells that appear to have migrated away from their clone of origin, we discarded all clones having less than 5 cells from the analysis.

**Figure 3.**
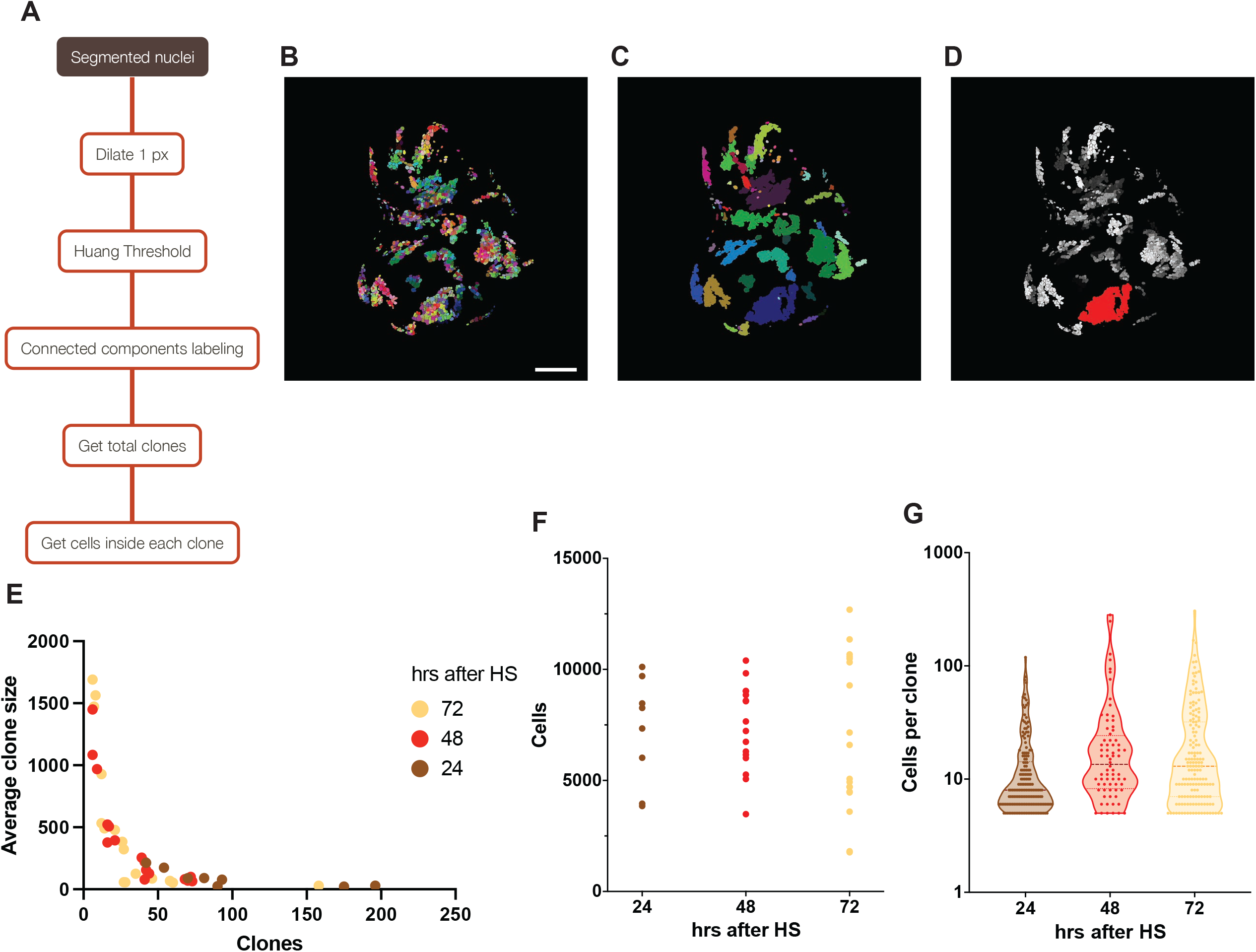
Individual clone statistics for nGFP-marked clones. A) Schematic of the clone analysis workflow, as performed on a watershed image processed by the nuclear counting workflow outlined in Fig. 1A. B) Max projection of a watershed wing disc with nRFP-tagged clones. C) Output image of the clone analysis workflow. Each clone is shown in a different color. D) Clone sizes on the imaginal disc shown on B. Clones of less than 5 cells have been excluded from the analysis. E) Number of clones per disc as measured in 42 wing discs, and the average clone size in cells for each of these discs. F) Total number of cells per disc for the three different heat-shock treatments. G) Number of cells per clone in three representative discs for each heat-shock treatment. Each dot of the violin plot represents one clone from that disc. For better representation, cells per clone are plotted as the logarithmic transform of the data on an anti-logarithmic Y-axes, as this is the preferred method of presenting logarithmic data on a violin plot without altering the distribution. Scale bar: 100 μm

To characterize the relationship between clone growth, time of clone induction, clone density, and clone fusion, we ran this workflow on clones tagged by act>CD2>Gal4, UAS-NLS::RFP generated by heat shock at 24, 48, or 72 hrs. before dissection in late third instar (Fig. 3E-G), and quantified the clones in a total of 42 discs. From the analysis, we obtained the number of clones per disc and the average number of RFP+ cells per clone. A long heat shock was used to achieve a high recombination rate, which resulted in a very dense clone population at 72 hrs after heat-shock (Fig. 3G). The quantification workflow revealed that the clone size is proportional to the time the discs had to develop after heat-shock as expected. In addition, is important to point out that larger and denser clones will show a greater tendency to merge into larger RFP+ regions, thus appearing to distort this relationship. This workflow might prove very useful in cell competition and other experiments with deleterious mutations to assess clone density, size, and the elimination of clonal cells.

We have presented here a rapid and accurate method to obtain data on cell number and volume of individual clones, providing a more comprehensive analysis that will improve upon and enrich currently employed analyses of clones. Validation of our method with GFP- and RFP-tagged clones, as well as DAPI-stained nuclei, makes us confident that this workflow can be applied to any reasonably robust nuclear fluorescent protein or marker.

### A trainable machine-learning algorithm to classify and count homozygous clones in a heterozygous background

Twin-spot analysis, in which induced mitotic recombination produces a clone homozygous for a marker and its sister clone that expresses no marker, in a heterozygous background, has been a staple in studies analyzing the effect of developmental processes affecting cell proliferation or cell death, such as cell competition or cancer. In those scenarios, one of the clones will carry a mutation that will alter growth, which after several rounds of cell division will result in clones in which the mutant clone grows to a different size compared to its twin-spot. Typically, these experiments have been scored by quantifying the total area of each twin-spot, as counting cells has proven extremely challenging. While clone area suffices in the most extreme cases, certain phenotypes can alter the clone area in a manner not proportional to the total number of cells (such as changes in polarity, adhesion, or mechanical tension), and perhaps more importantly, even in control discs, area poorly correlates with cell number, as shown above. Therefore, a method to accurately count cells in twin-spots would greatly benefit this kind of analysis.

Adaptation of our nuclear counting script to the analysis of twin-spots requires additional steps in order to automatically distinguish homozygous cells from the heterozygous background. To exclude cells that do not belong to the homozygous clones, we first tried to remove the heterozygous cells from the counting workflow with a simple intensity segmentation. However, nuclear intensity varies in different areas of the wing disc, such as the hinge and the pouch, or the folds on the dorsoventral boundary (Fig. 4A). Therefore, a simple segmentation either picks up many false positives or ignores homozygous clones in dimmer areas of the disc. We therefore developed an algorithm that could consider characteristics of the local neighborhood when classifying cells and clones (See Methods; Suppl. Fig. 5). We used Ilastik(Sommer *et al*., 2011; Berg *et al*., 2019), a comprehensive suite of workflows powered by trainable machine learning, to train a Random Forest machine learning algorithm to automatically classify clones according to their surrounding context.

**Figure 4.**
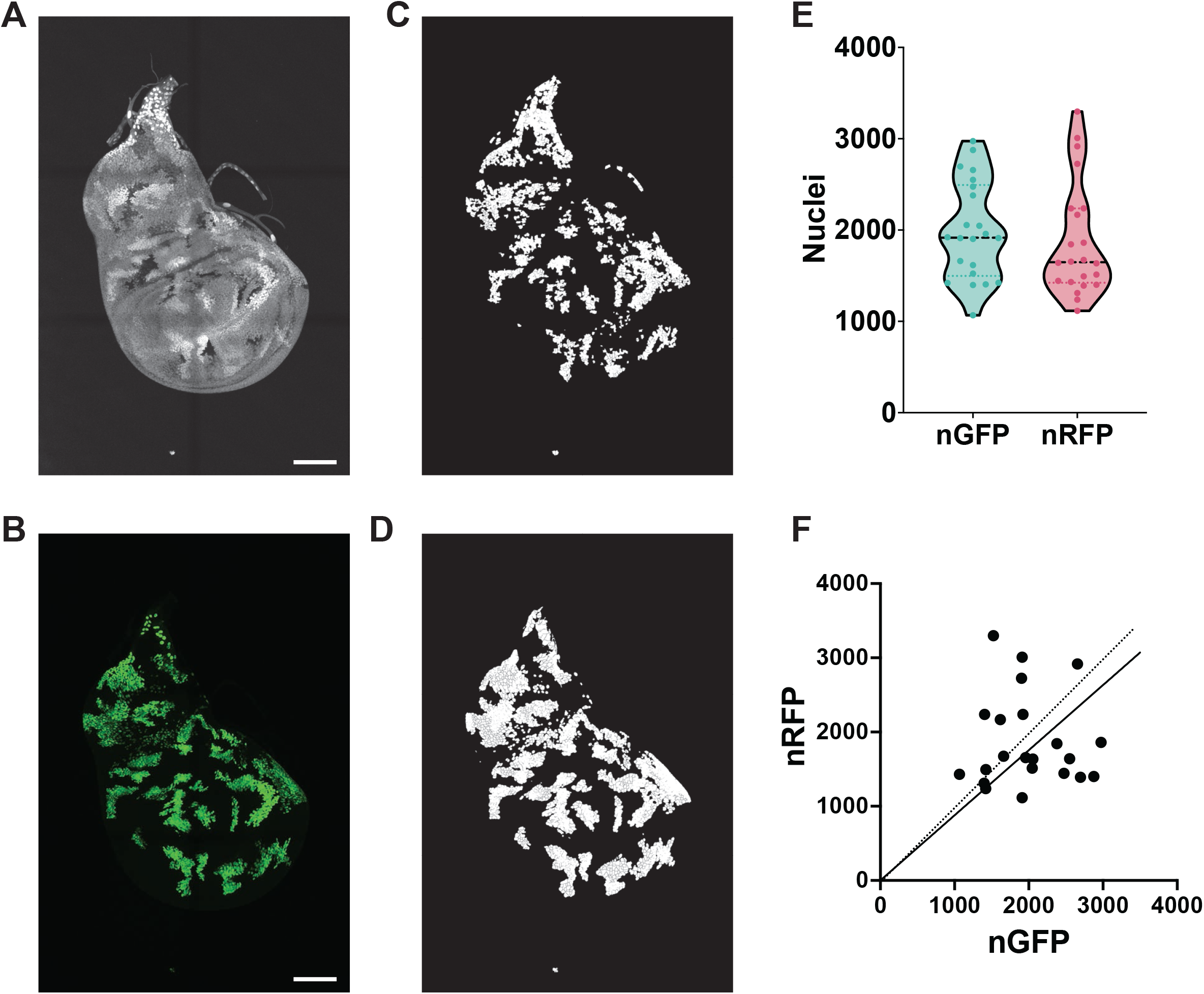
Segmentation and counting of twin spots. A) tub-nRFP heterozygous and homozygous nuclei of a representative disc used for the training of the segmentation algorithm. B) UAS-nGFP clones; here, nGFP is expressed only in the homozygous tub-nRFP twin-spots. C) Ilastik’s segmentation of homozygous tub-nRFP cells from the disc shown in A. D) Segmentation of the UAS-nGFP-marked cells from the disc shown in B. E) Overall counts on 22 discs containing homozygous twin spots marked with nRFP. Each data point of the violin plot shows the total nGFP and nRFP cell counts per disc, median and quartiles are indicated. F) Correlation plot of nGFP vs nRFP counts for each analyzed discs, showing the disparity for nuclei counted with our analysis. The solid line represents the best linear fit for all data points and the dotted line represents the perfect 1:1 fit. N = 22 discs. Scale bar: 100 μm

We trained Ilastik’s Object classification workflow (for more information on the different Ilastik workflows, see (Berg *et al*., 2019)) to discriminate between heterozygous and homozygous NLS::RFP nuclei. For the training, we used flies harboring a genetic system that creates twin-spot clones marked with NLS::RFP that will also activate the expression of NLS::GFP (FIG 4A-B; see methods). As the homozygous RFP clones are the only cells in the disc that express GFP, the GFP channel serves as an internal control that allows us to easily visualize which cells are part of homozygous RFP clones, a distinction that can challenge the observer due to disc contours and regional intensity variation. We trained the segmentation algorithm using 9 discs to account for variable overall image intensity, disc size, clone size, and orientation. To validate our RFP segmentation and counts, we also counted the cells expressing GFP as a control. If trained correctly, our pipeline should count approximately the same number of GFP and RFP cells per disc. After manually training the algorithm as detailed in the methods section (Suppl. Fig. 6), we applied it to 22 discs, including the 9 used for training and 13 not used for the training. We counted RFP nuclei on the segmented homozygous clones (Fig. 4C), and compared the cell counts to the GFP clones (Fig. 4D).

Side by side comparison of the RFP counts and GFP controls (Fig. 4E) demonstrated good accuracy, with the average values deviating by less than 10% (1992 ± 538 GFP nuclei vs 1874 ± 620 RFP nuclei). Although similar when analyzed as a group, a pairwise comparison between nRFP and GFP counts in individual discs showed a surprising degree of variation (Fig. 4F). We initially presumed that the difference was attributable to occasional errors introduced by the segmentation and classification workflow, which might misidentify a fraction of the RFP cells. However, visual inspection of the images revealed individual clones in most discs that had different numbers of GFP and RFP labelled nuclei (Fig. 4, A-B and Suppl. Fig. 6). Manual counting of some of these discordant clones showed that the classification and counting of RFP cells was very accurate, showing little or no misclassified nuclei (Fig. 4 A,C and Suppl. Fig. 6). Therefore, a large fraction of the difference between GFP and RFP counts can be attributed to differences in the behavior of the transgenes driving GFP and RFP expression (UAS-NLS::GFP versus tub-NLS::RFP), rather than errors in the RFP segmentation. Because our workflow facilitates the rapid and easy counting of large numbers of clones and discs, we suggest that a modest level of inaccuracy in individual counts will be more than sufficiently compensated by the statistical power achievable with our method.

### Conclusions

The three user-friendly cell count pipelines presented here can be applied to any image without having to perform specific alterations to the pipeline. We have applied our pipelines to DAPI-stained cells and UAS-NLS::GFP, UAS-NLS::RFP, or tub-nRFP marked cells. Nevertheless, as the workflows are applied to greyscale images, they can most likely be used with any other nuclear dye or fluorescent protein, provided that they are tagged with a NLS to be localized to the nucleus.

Typically, clones have been analyzed by a) presence/absence of clones (Moreno and Basler, 2004); b) number of clones(Martín *et al*., 2009; Merino *et al*., 2015) ; or c) clone size (de la Cova *et al*., 2004; Meyer *et al*., 2014; de Vreede *et al*., 2022). While often sufficient for a qualitative assessment of the resulting effect, the pipelines described here, created by combining classic methods, such as watershedding (Soille and Vincent, 1990) with machine learning by trainable algorithms (Sommer *et al*., 2011; Berg *et al*., 2019), provide tools to go beyond these simple measurement outputs to derive more nuanced and quantitative descriptions of the behavior of wing disc clones and cells.

In clonal analyses, acquiring cell counts in addition to clone volumes and areas will provide more richly featured descriptions that will be informative when analyzing the effects of genes that could alter cell shape or size, such as structural proteins or genes related to tissue integrity (Lecuit *et al*., 2011; Borghi *et al*., 2012; Fulford and McNeill, 2020). In these cases, area measurement alone would result in misleading interpretation of results, whereas a more comprehensive analysis including measuring direct cell counts and clone volumes will provide for a more precise interpretation of mutation effects (Fig. 1F, Suppl. Fig. 3). The tools presented here will pave the way toward easily achieving comprehensive quantitative analyses.

## Methods

### Fly stocks and husbandry

The following fly lines were used:

- To count cells by DAPI staining: OreR (WT flies)
- To analyze GFP or RFP clones over a WT background: *y*^*1*^ *w*^*1118*^ *hsp-Flp act-Scer/Gal4 Scer/UAS-NLS::GFP;* P{neoFRT}42D *tub-Scer/Gal*^*80*^ / P{neoFRT}42D and *y*^*1*^ *w*^*1118*^ *hsp-Flp act>CD2>Scer/Gal4 Scer/UAS-NLS::RFP*
- To count homozygous GFP and RFP labeled cells and unlabeled cells in twin-spot analyses: *y*^*1*^ *w*^*1118*^ *hsp-Flp act-Scer/Gal4 Scer/UAS-NLS::GFP;* P{neoFRT}42D *tub-Scer/Gal*^*80*^ *tub-NLS::RFP* / P{neoFRT}42D

Flies were maintained in standard fly food in a temperature-controlled incubator at 25ºC. Egg collections were performed over a timespan of 24 hours. Vials with eggs were kept at 25ºC until dissected or heat-shocked for clone generation.

Late first instar and early second instar larvae were heat-shocked for 15 min, 72 hours before dissections. Heat-shocked larvae were kept at 25ºC until they were dissected. Wandering third-instar larvae were harvested for analysis.

### Generation of clones and sample preparation

Wing imaginal discs were dissected by inverting the larva and removing the fat body and digestive tissues (mouth hooks, salivary glands and intestine) from the anterior half.

Inverted larvae were fixed in 4% Paraformaldehyde (PFA) in phosphate buffered saline (PBS) + 0.02% Triton X–100 (PBS-T) for 30 min at room temperature (RT). Discs were then washed 3 times with PBS-T and then stained with DAPI in PBS-T for 5 min. Discs were then washed 3 times with PBS, and then carefully extracted from the inverted head and mounted in Vectashield. Special care was taken to remove all the tracheal tissue from the wing disc as its autofluorescence would interfere with subsequent analysis.

### Image acquisition

Images of whole discs were taken with a Leica SP8 system equipped with a White Light Laser and HyD(R) detectors. A 40x, NA of 1.5 Leica objective and 1.51 refractive index immersion oil were used. Channels were acquired separately to avoid cross-talk, and the laser power and sensor gain were optimized to avoid histogram saturation. Images were taken in 8 bits, at 1024×1024 resolution, and a pinhole of 1 Airy Unit (AU) and acquired using a Z scan with a step size of 0.3 μm and a variable total Z depth to encompass all the disc proper cells and as few peripodial cells as possible. Leica’s software was then used to stitch a tilescan of different fields of view to image the entire disc. Tilescans were imported into Fiji (fiji.sc, (Schindelin *et al*., 2012)). Before analysis, any leg and haltere discs in the cases where they remained attached to the wing disc were we cropped out from the images. Apical and basal slices of the Z stack were also removed if needed to get rid of most or all the peripodial cells. While preferred, this step is not critical, as peripodial cells normally account for less than 5% of the wing disc cells [Shubiger 2004] and the acquisition parameters of our Z stacks already excluded most of the peripodial cells. Images were then analyzed using the nuclear counting workflows detailed below.

### Workflows

Tiff tilescans were processed using Fiji macros that perform functions readily available in the Fiji ImageJ distribution and from the MorphoLibJ library (Legland *et al*., 2016), which is included in the Fiji package. For more information about installing and using the MorphoLibJ library, see https://imagej.net/plugins/morpholibj. Each of the pipelines described here is coded into a custom ImageJ macro for a single-click analysis. Although the performed steps for each analysis are described below, we recommend using the custom macros for a user-friendly process. Macros can be downloaded from GitHub (LINK) and are available upon request.

### Wing disc total nuclei counts

The nuclear count workflow is summarized in Fig. 1A. Tiff images were first processed through a 2px 3D Gaussian Blur to remove noise and allow for optimal segmentation through watershedding. Images were then inverted prior to segmentation of nuclei by watershed. Inverted images were processed using MorphoLibJ’s Classic Watershed to segment all nuclei, without mask, and including intensities from one point below the image minimum intensity to 250. The watershed nuclei are then counted using MorphoLibJ’s Connected Components Labeling, using 26-point connectivity and a 16-bit output result. Residual noise is removed by filtering any object smaller than 10 voxels in volume using MorphoLibJ’s Label Size Filtering and then the labels were re-assigned by using the MorphoLibJ plugin Remap Labels. By using this pipeline, each segmented nucleus is assigned an individual value on the histogram of a 16-bit image file, starting with 1 and up to 65536. The total number of labels can be manually checked by looking at the image histogram, but to automatically obtain the total nuclear counts via macro, we coded it to measure the highest intensity in the whole image stack, which corresponds to the last label in the image (hence giving us the total number of nuclei). To do so, we measure the Max. intensity on the stack’s Max. Projection. The final numbers are outputted to Fiji’s Result table and, if using our macro, the log window.

To perform the workflow described above with our Fiji macro, open a single channel image file containing the whole disc with stained nuclei and use the command Plugins>Macro>Run… from Fiji to select and apply the macro “DAPI_counts_with_MorphoLibJ.ijm” to the open image. To batch analyze several single-channel images in a folder, use the command Process>Batch>Macro… in Fiji. Then choose the folder containing the files as input and copy the macro code into the text window provided. Additional information on batch processing can be found at https://imagej.net/scripting/batch. Alternatively, we provide a batch processing macro that allows through a simple user interface to choose a folder with files and one of our macros to apply it to all files in the chosen folder. To use it, simply execute the “Batch_process_folder.ijm” macro.

### Clone analysis on GFP and RFP nuclei

GFP- and RFP-labeled nuclei in a WT background were counted by first using the nuclear count analysis outlined above. Then, the Output Labels were processed by the following workflow, as outlined in figure 3A, to obtain the number of clones and cells per clone:

To create a binary mask to segment the individual clones, Nuclei Labels were first dilated 1 px using MorphoLibJ’s Dilate Labels plugin, then converted into a binary mask using Huang’s Thresholding. The binary mask was then run through MorphoLibJ’s Connected Components Labeling using the same parameters as for DAPI nuclei. Then every label smaller than 100 voxel was removed with MorphoLibJ Label Size Filtering. Finally, the total number of clones was obtained and outputted as with the nuclear counts, using Remap Labels followed by measuring the last labeled clone as the highest intensity in a Max. Projection of the image. This workflow provides the total number of clones in the imaginal disc. To calculate each clone size, each clone is isolated using MorphoLibJ’s Select Label(s) plugin. The resulting image is multiplied to the segmented Nuclei Labels (the Output Labels obtained through the nuclear counting workflow) using Fiji’s Calculator Plus. This step provides a new image including only the cells contained within the clone. Finally, those cells are counted using Connected Components Labeling as previously done for the whole disc.

As for DAPI counts, a convenient ImageJ macro is provided for a single-click analysis. On a single channel image containing the fluorescently tagged clones, apply the macro “GFP_counts_with_MorphoLibJ.ijm”. For batch analysis, follow the same steps as for the DAPI counts to apply the code into a folder containing single-channel images.

### Volume computation of whole discs and of labeled clones

To quantify the disc and clone volumes, we used the segmented image obtained from the nuclei counts and applied the MorphoLibJ’s plugin Analyze Regions 3D to measure each label’s volume on the image. After the watershed and nuclear segmentation, the labels are expanded so that only 1 px separate them from each other (Figs. 1-C’ and 2B).

Therefore, the segmented images serve as an accurate readout of the total cell volume. Analyze Regions 3D provides a table with all the cell volumes, which are then added to obtain the total volume for either the whole disc (in the DAPI-stained total cell counts) or the fluorescently labeled clones. This step is included in the nuclear counting and clone analysis ImageJ macros, as it is very efficient and adds very little computational time to the macros.

### Machine-Learning segmentation of twin-spot clones

In twin-spot analyses, a homozygously labeled clone and its unlabeled twin-spot are produced for each recombination event. The homozygously labeled clone must be distinguished from the heterozygously labeled unrecombined cells, presenting a challenge due to variable intensity across the disc. The following method (Suppl. Fig. 5A) accurately accomplished this step.

ML-based image segmentation was performed with Ilastik (https://www.ilastik.org, (Sommer *et al*., 2011; Berg *et al*., 2019)), by using the Object classification workflow. The trained algorithm is available upon request. It can either be opened and used from Ilastik or called using ImageJ’s Ilastik plugin (To download, check https://www.ilastik.org). However, due to the sensitivity of the Object Classification workflow and the modest time required to train the algorithm, we suggest training a new Object Classification workflow *de novo* for datasets the user wishes to evaluate following the steps described below.

Tiff images from imaginal discs were first processed for the intensity-based object classification workflow. Intensity was normalized using Fiji’s Enhance Contrast processing tool, with a 0.1% of saturated pixels. The normalized files were exported from Fiji as HDF5 files (Suppl. Fig. 5B). To segment the nuclei for the Ilastik classification, each file was processed via watershed following the same pipeline used for counting DAPI nuclei. The watershed was converted into a binary mask using Fiji’s Convert to Mask command and then exported as HDF5 (Suppl. Fig. 5C). Both the normalized image and the segmentation mask were imported into the Ilastik’s Object Classification to perform the training. The algorithm was manually trained using 9 imaginal discs.

To classify the homozygous RFP clones, only the intensity features were selected. The neighborhood of each pixel was set to 50 px (Suppl. Fig. 5D). The algorithm was trained by manually classifying 10-50 cells on each disc as heterozygous or homozygous (Suppl. Fig 6E). Using the Live Preview on Ilastik we fine-tuned the classification by inspecting the automatic classification on the training dataset and manually correcting wrongly classified cells. When the algorithm was validated on the training dataset we applied it to the whole dataset of 22 imaginal discs. The segmented images were exported as Object Predictions (Suppl. Fig. 6F) and imported into Fiji to count the nuclei classified by Ilastik as homozygous RFP. The imported file, which include both heterozygous and homozygous nuclei, was then converted to a binary mask using a Default threshold of 2 to remove all cells classified by Ilastik as heterozygous. Nuclei were then counted using the same pipeline used for DAPI and GFP nuclei, starting from MorphoLibJ’s Connected Components Labeling.

In twin-spot analysis, the unlabeled twin-spots can be estimated by counting the total number of fluorescently-labeled nuclei (homozygous plus heterozygous) and subtracting it from the total DAPI nuclear counts. This calculation will provide with the number of DAPI-stained nuclei that are not marked by the fluorescent protein label.

As with the other workflows, the macro script allows one to perform the whole analysis in Fiji without the need for user interaction. Just execute the “Twin-spot_counts_with_MorphoLibJ.ijm” macro on a single-channel image containing the twin-spots labeled with a nuclear fluorophore. Alternatively, the macro can operate in batch mode by applying this code to a folder containing the images as described for the DAPI nuclei count.

## Acknowledgements

We would like to thank members of the Axelrod lab for fruitful discussions. Stocks obtained from the Bloomington Drosophila Stock Center (NIH P40OD018537) were used in this study.

This work was supported by the National Institutes of Health grant R35 GM131914 to JDA (https://www.nigms.nih.gov) and the Swiss National Science Foundation (<u>P400PB_199258</u>) to PSB. The funders had no role in study design, data collection and analysis, decision to publish, or preparation of the manuscript.

## Conflict of interest

The authors declare that the research was conducted in the absence of any commercial or financial relationships that could be construed as a potential conflict of interest.

## Supplementary figure legends

**Supplementary figure 1.**
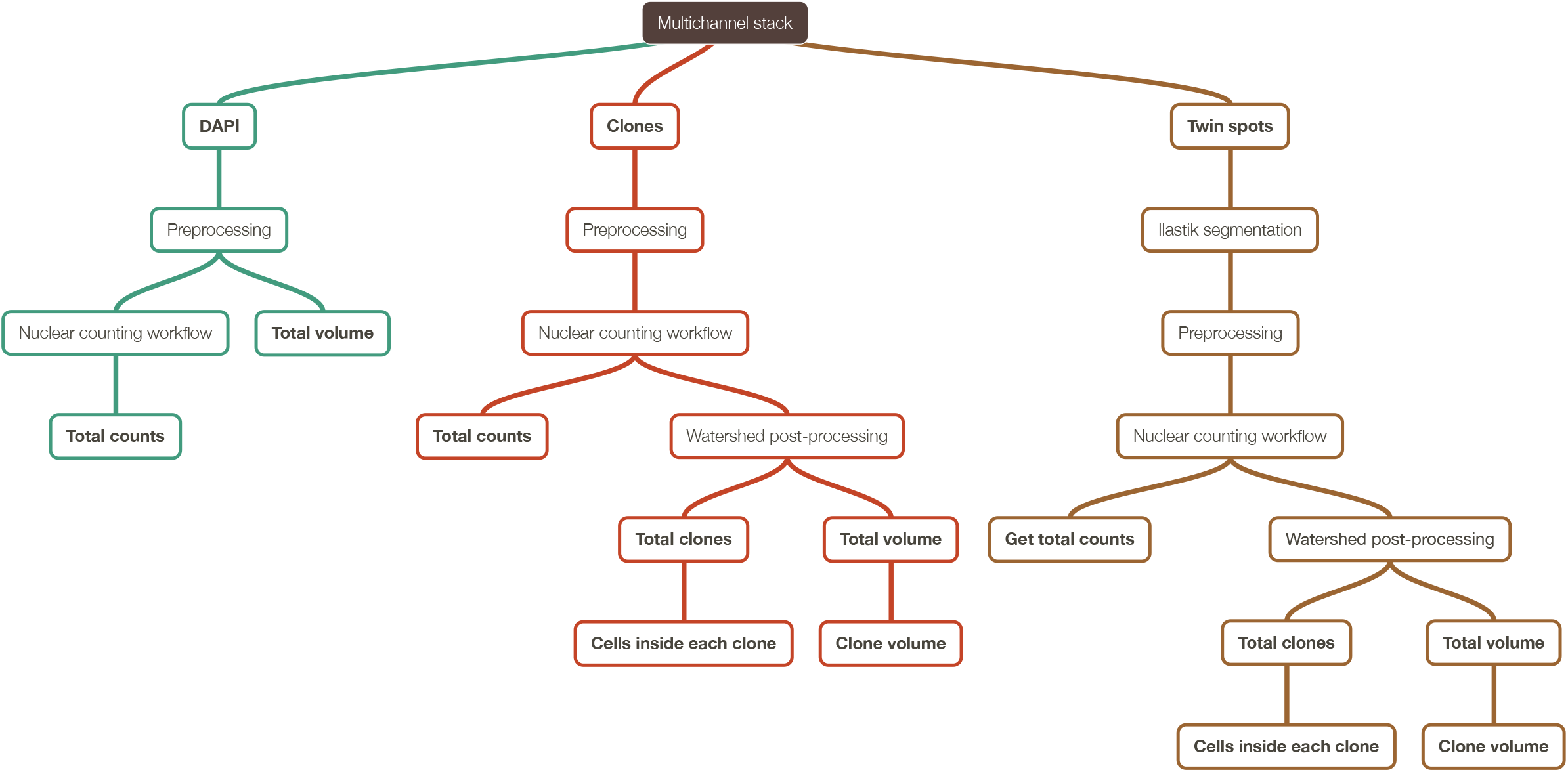
Summary of the three nuclear counting wokflows. Overall workflow for classification and counting of wing disc nuclei, summarizing the measurements obtained with our method.

**Supplementary figure 2.**
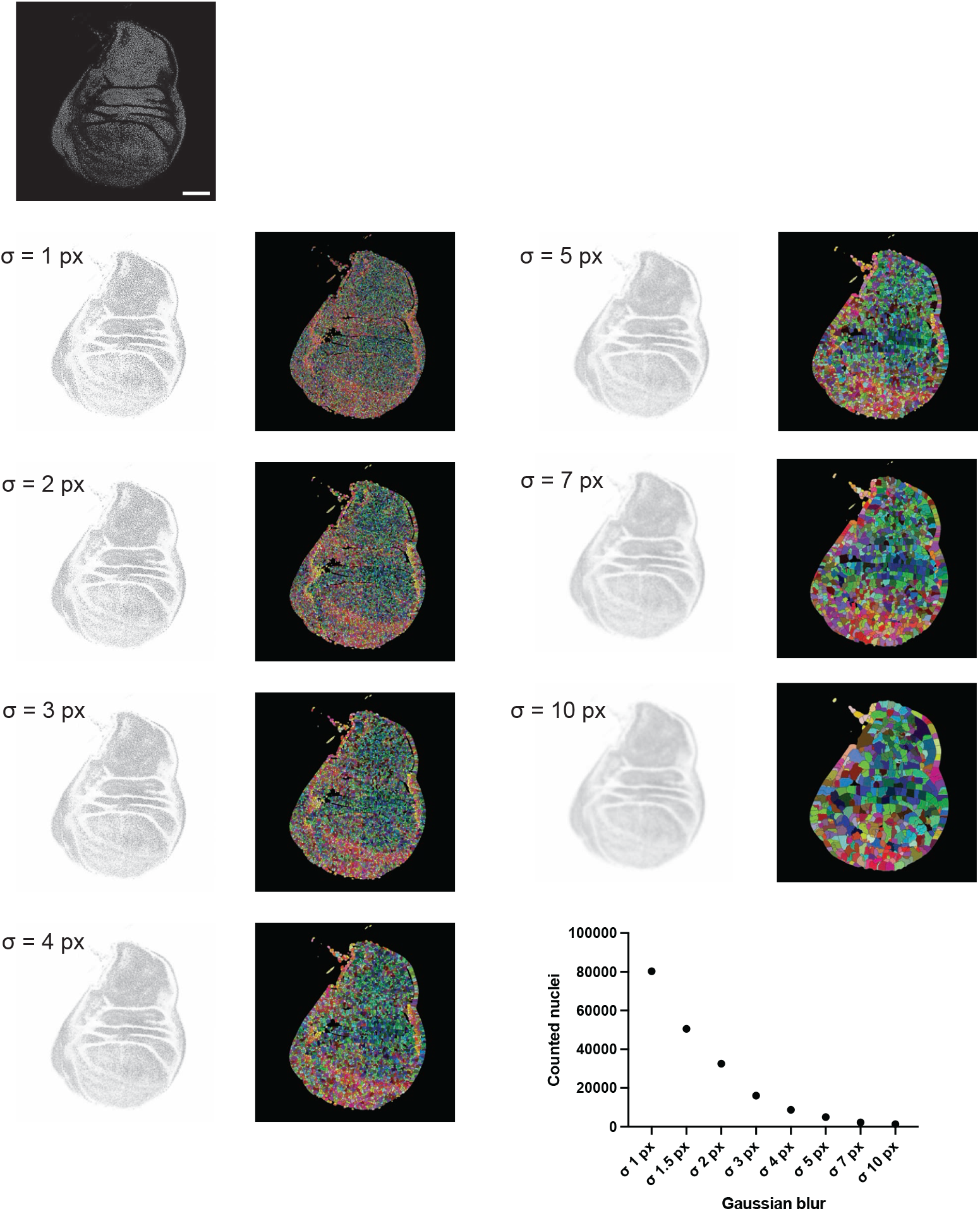
Effect of gaussian blur on counting efficiency. Images show the overall look of the disc and the resulting watershed for different sigma values of the 3D Gaussian blur on a representative DAPI-stained disc. The graph shows the counts on the example disc for the different sigma values. Scale bar: 100 μm

**Supplementary figure 3.**
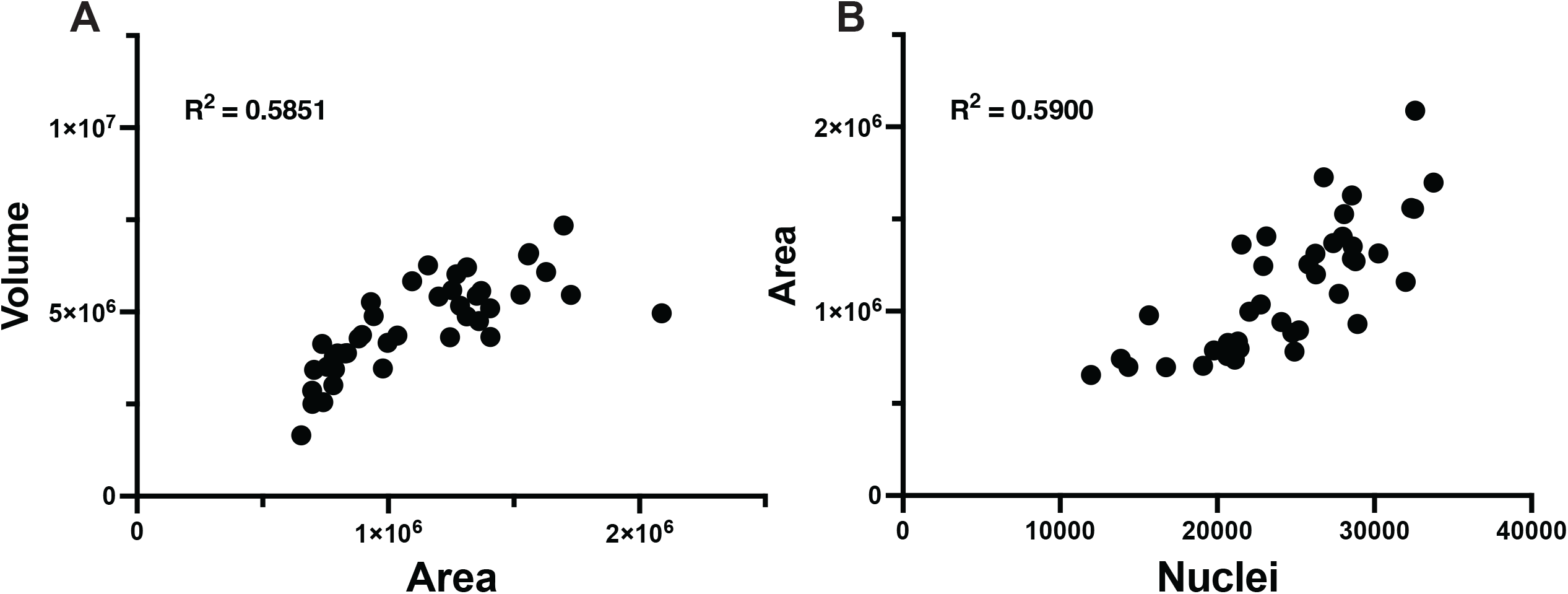
Quantifying disc size using area measurements. A) Correlation plot showing the results of measuring disc size by area vs volume. The R^2^ of 0.7212 indicates that the measures don’t correlate significantly. B) Correlation between area and nuclear density. The number of nuclei does not correlate with the area the wing disc covers (R^2^ of 0.5909).

**Supplementary figure 4.**
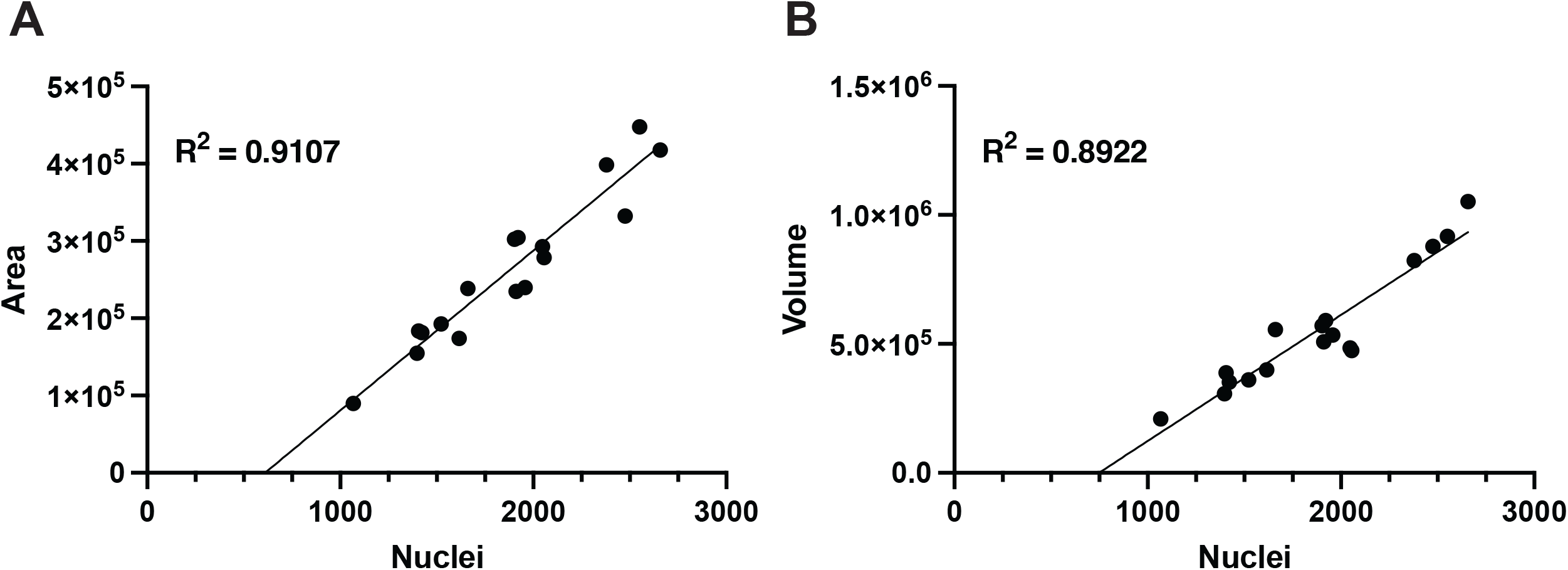
Correlation between area, volume and nuclear counts in low-density clones. A) Correlation between area and nuclear counts for the 17 discs in the NLS::GFP dataset used for clone counts (R^2^ of 0.9107). Correlation between volume and nuclear counts for the same dataset (R^2^ of 0.8922).

**Supplementary figure 5.**
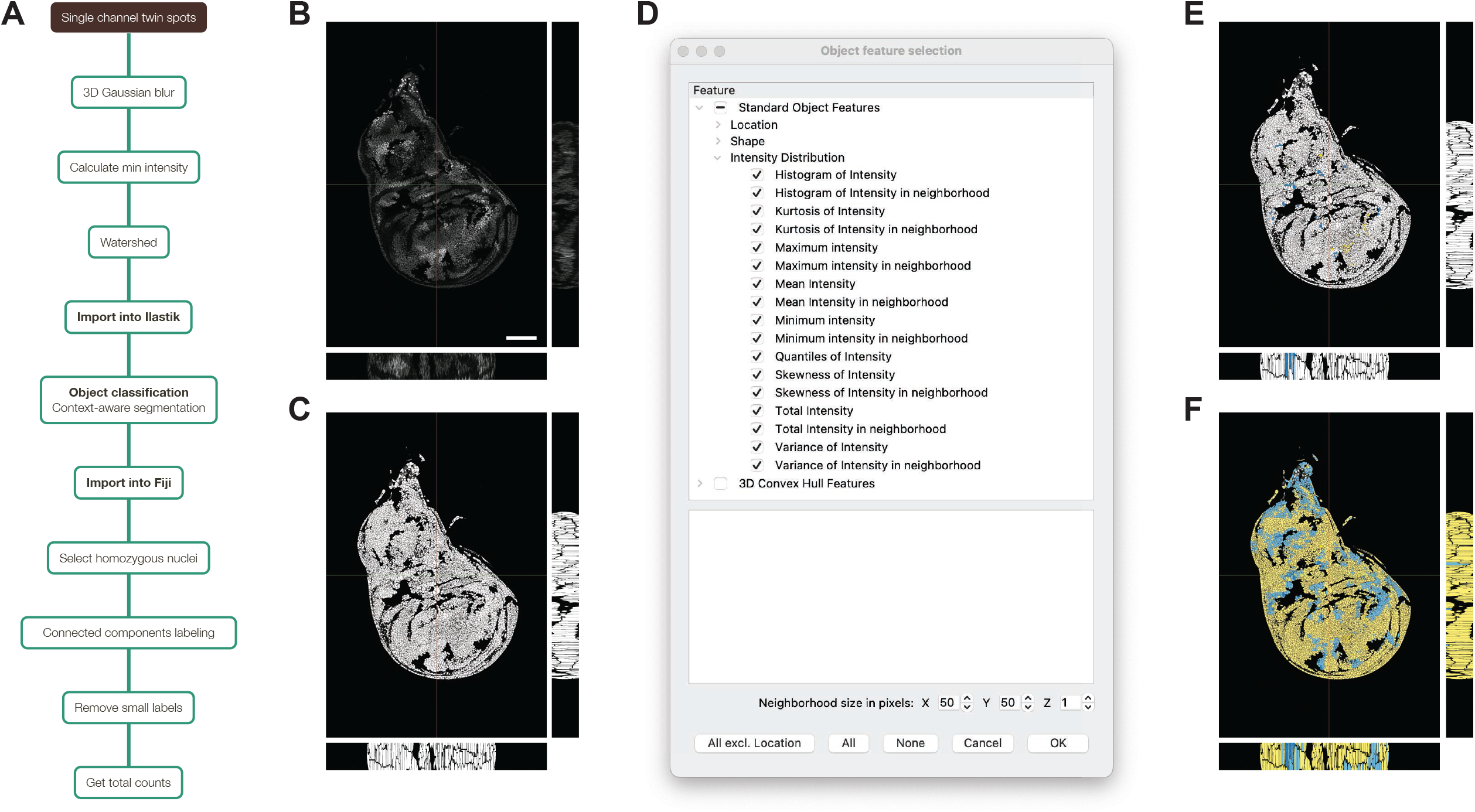
Twin-spot classification overview. A) Schematic of the segmentation of homozygous twin-spots using Fiji and Ilastik’s Object classification. B) Raw input image passed on to Ilastik for the clone classification. C) Segmentation image used by Ilastik to classify each nucleus. D) Computation parameters used to classify each nucleus as heterozygous or homozygous according to its intensity and the surrounding 50×50 pixels. E) Training of the object classification algorithm, showing a few nuclei manually classified as heterozygous (yellow) or homozygous (blue). F) Prediction for every nRFP nucleus as heterozygous (yellow) or homozygous (blue) in the wing disc based on the trained algorithm. All wing disc images show the central slice of the Z stack, with the orthogonal views corresponding to the sections highlighted in green and red. Scale bar: 100 μm

**Supplementary figure 6.**
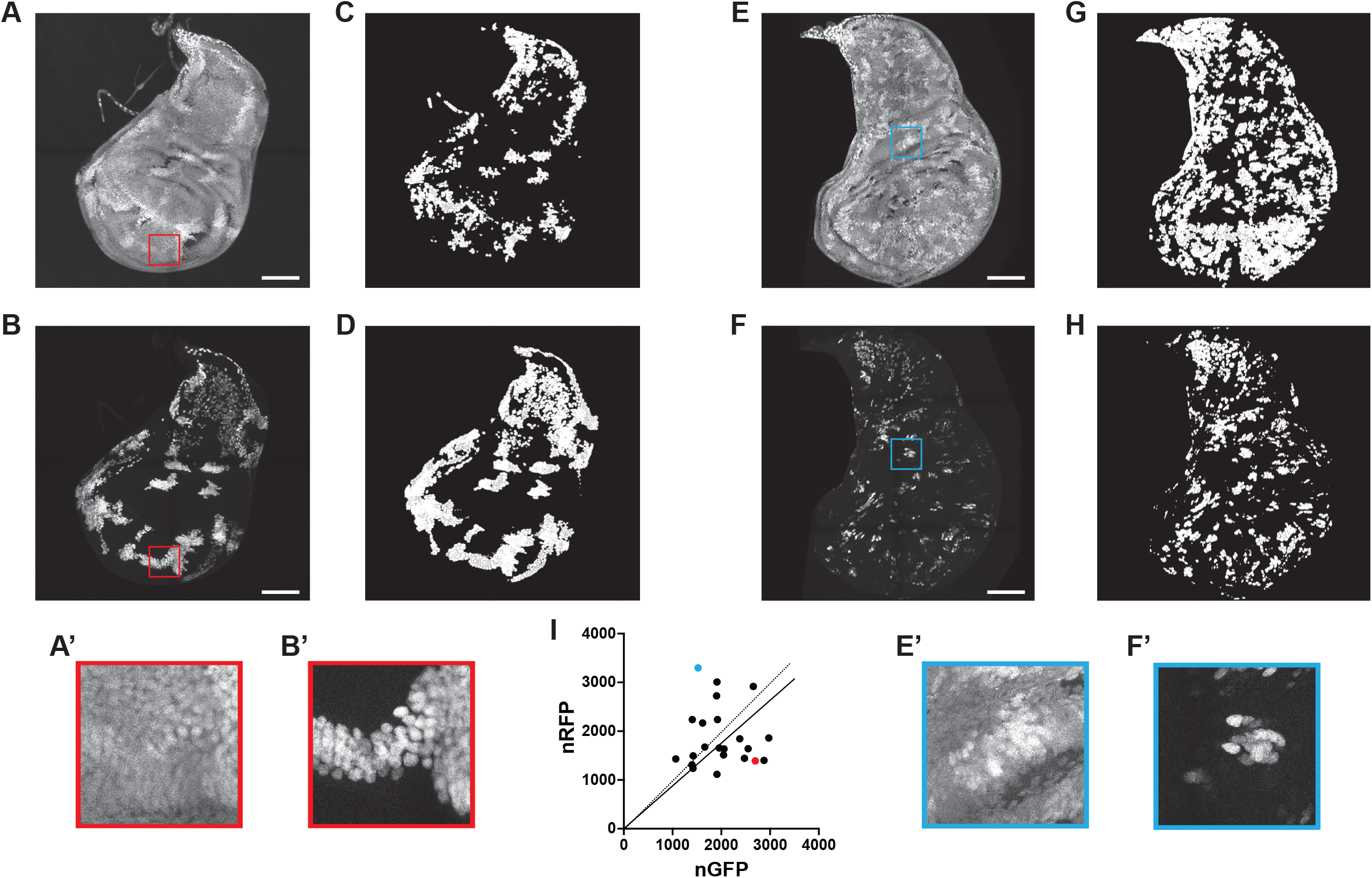
Differential expression of tub-nRFP and UAS-nGFP in twin spot clones. A-D) Representative disc in which GFP nuclei are more numerous than RFP nuclei. A’, B’) Closeup of the section highlighted in red in A and B. E-H) Representative disc in which GFP nuclei are less abundant than RFP nuclei. E’, F’) Closeup of the section highlighted in blue in E and F. A, E) nRFP twin-spot nuclei. B, F) Homozygous nRFP twin-spot nuclei classified by Fiji and Ilastik as summarized in the methods. C, G) GFP nuclei. D, H) Segmentation of GFP nuclei. I) Correlation plot comparing nRFP and nGFP in 22 discs. The representative disc in (A) is indicated by the red dot and in (E) by the blue dot. Scale bar: 100 μm

